# Ribosome profiling in *Streptococcus pneumoniae* reveals the role of methylation of 23S rRNA nucleotide G748 on ribosome stalling

**DOI:** 10.1101/2020.12.21.423859

**Authors:** Tatsuma Shoji, Akiko Takaya, Yoko Kusuya, Hiroki Takahashi, Hiroto Kawashima

## Abstract

2.

**(1) Background:** Many nucleotides in 23S rRNA are methylated post-transcriptionally by methyltransferases and cluster around the peptidyltransferase center (PTC) and the nascent peptidyl exit tunnel (NPET) located in 50S subunit of 70S ribosome. Biochemical interactions between a nascent peptide and the tunnel may stall ribosome movement and affect expression levels of the protein. However, no studies have shown a role for NPET on ribosome stalling using an NPET mutant.

**(2) Results:** A ribosome profiling assay in *Streptococcus pneumoniae* demonstrates for the first time that an NPET mutant exhibits completely different ribosome occupancy compared to wild-type. We demonstrate, using RNA footprinting, that changes in ribosome occupancy correlate with changes in ribosome stalling. Further, statistical analysis shows that short peptide sequences that cause ribosome stalling are species-specific and evolutionarily selected. NPET structure is required to realize these specie-specific ribosome stalling.

**(3) Conclusions:** Results support the role of NPET on ribosome stalling. NPET structure is required to realize the species-specific and evolutionary conserved ribosome stalling. These findings clarify the role of NPET structure on the translation process.

## 4. Background

Endogenous rRNA modifying enzymes methylate or pseudouridylate specific rRNA nucleotides at functionally important regions in the ribosome, such as the peptidyl transferase center (PTC) [1]. Approximately one-third of modified residues of 23S rRNA are clustered around the nascent peptide exit tunnel (NPET) [2]. While the role of rRNA modification remains unclear, though it is generally believed that they have fine-tune functions of the ribosome in translation [3], especially under the stress conditions [4], via the biochemical interactions between the nascent peptide and tunnel [5,6]. These interactions may stall ribosome movement and thus affect the expression level of the protein [6]. However, no studies have shown the role of NPET in ribosome stalling using an NPET mutant.

Some modifications of NPET are important for determining antibiotic resistance or susceptibility [7,8]. While the role of the methylation at G748 (m^1^G748) in *Streptococcus pneumoniae* remains unclear, we previously showed that inactivation of the methyltransferase RlmA^II^, which methylates the N-1 position of nucleotide G748 located near the PTC, results in increased resistance to telithromycin (TEL) in *erm*(B)-carrying *S. pneumoniae* [7].

We explored the role of NPET structure in translation initially by establishing the ribosome profiling assay in *S. pneumoniae* and investigating ribosomal distribution in both wild-type and RlmA^II^-deficient *S. pneumoniae*. Subsequent analysis showed that m^1^G748 is responsible for ribosome stalling and plays a role in species specificity.

## 5. Results

### (1) The loss of N-1 methylation at G748 greatly changes the distribution of ribosomes

We constructed two *S. pneumoniae* mutant strains, Sp284 and Sp379 to investigate the role of NPET. Sp284 is a RlmA^II^-disrupted mutant harboring pTKY1111 encoding *tlrB* of the S1 strain [7]. Sp379 is a RlmA^II^-disrupted mutant harboring pTKY1127 encoding *tlrB* of Sp44 [7]. The latter strains display no methyltransferase activity because of the C23R mutation [7]. The difference between Sp284 and Sp379 is only the capability to methylate G748.

We also performed a deep sequencing-based ribosome profiling analysis in Sp284 and Sp379. Ribosome profiling captures a global snapshot of ribosome positioning and density on template mRNAs with single-nucleotide resolution. The *erm*(B) operon, where ribosomes stall at the *ermBL* region [9], was selected, for example, to examine the distribution of ribosomes (Fig. 1A). Ribosome position and density in the *erm*(B) operon in Sp379 was completely different from positions and density in Sp284 (Fig.1A).

**Fig 1.**
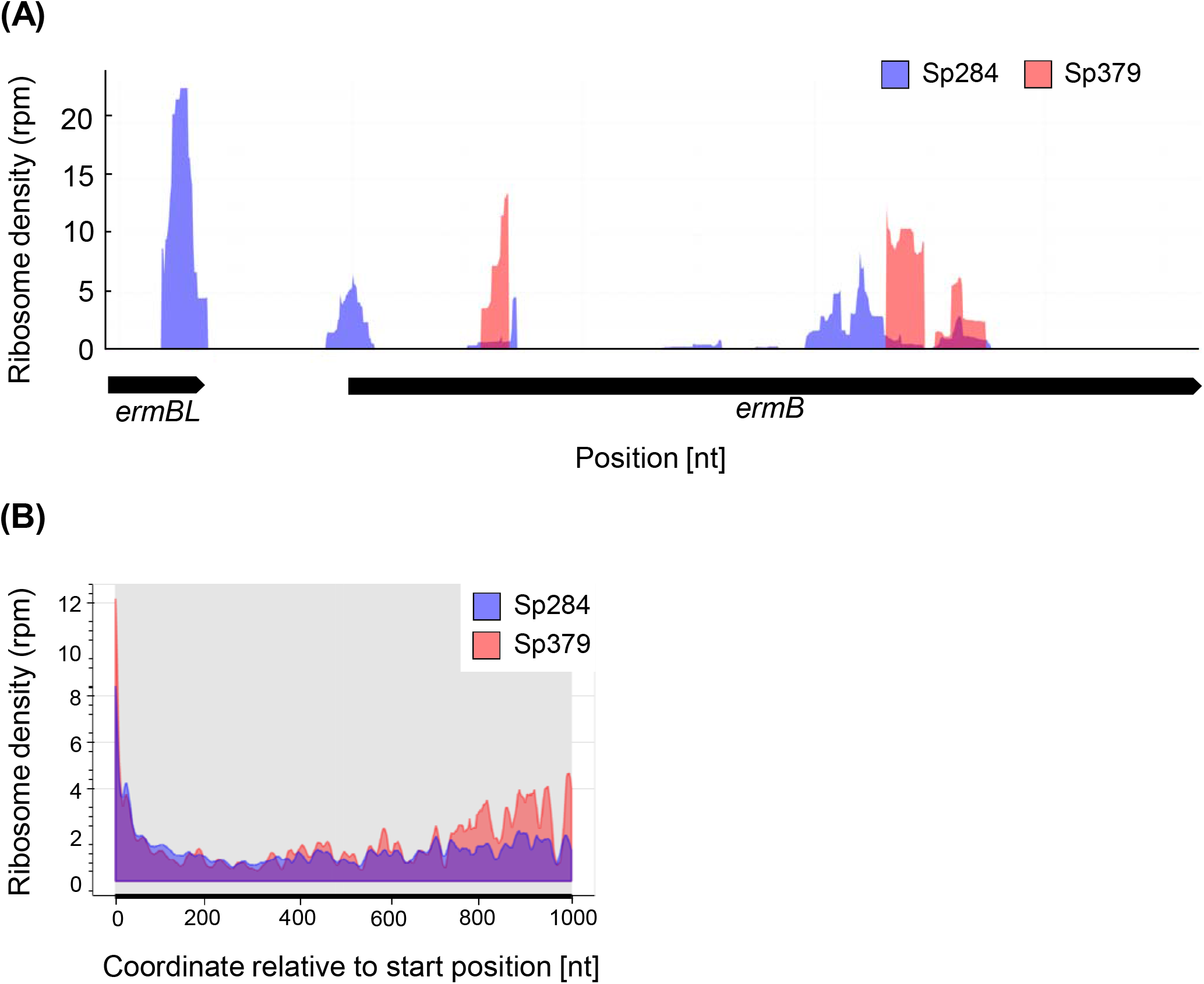
Differences in ribosome density (RD) profiles between Sp284 and Sp379. (A) RD in *erm*(B) in Sp284 or Sp379. (B) Metagene profile of RD in Sp284 or Sp379. RD across all ORFs was aligned relative to the start position.

The difference in the ribosome occupancy between two strains in *erm*(B) operon led us to speculate a general role of m^1^G748 in the translation process. Ribosome density across all ORFs in Sp379 was higher than in Sp284, especially around the latter region, indicating an important role of m^1^G748 in the translation process, with differences in ribosome position and density across all open reading frames (ORFs) being assessed using a metagene analysis (Fig. 1B).

### (2) m^1^G748 affects ribosome stalling

A high number of ribosome-protected footprints (RPFs) mapping to the transcriptome (mRNA-seq) at a unique position is indicative of ribosome stalling [10]. Thus, observed global changes imply a role for m^1^G748 in ribosome stalling. We tested this hypothesis by first constructing *erm*(B) operon-overexpression *S. pneumoniae* strains Sp380 and Sp382 from Sp284 and Sp379 respectively, to clarify differences between Sp284 and Sp379 (see Tables 1 and 2 for details). We then probed ribosomes of Sp380 and Sp382 before and after extracting total RNA with dimethyl sulfate.

**Table 1.** Bacterial strains.

| Strain | Relevant characteristics | Reference of source |
| --- | --- | --- |
| *Streptococcus pneumoniae* |  |  |
| S1 | TEL resistance clinical isolate | [7] |
| Sp36 | TEL-resistant mutants of S1 isolated from 8<br>μg/mL TEL-containing BHI-Y agar plates | [7] |
| Sp274 | $\Delta tlrB::aad(9)$ in S1 | [7] |
| Sp284 | Sp274 harboring pTKY1111 | This study |
| Sp379 | Sp274 harboring pTKY1127 | This study |
| Sp380 | S1 harboring pTKY1041 | This study |
| Sp382 | Sp274 harboring pTKY1041 | This study |

**Table 2.**
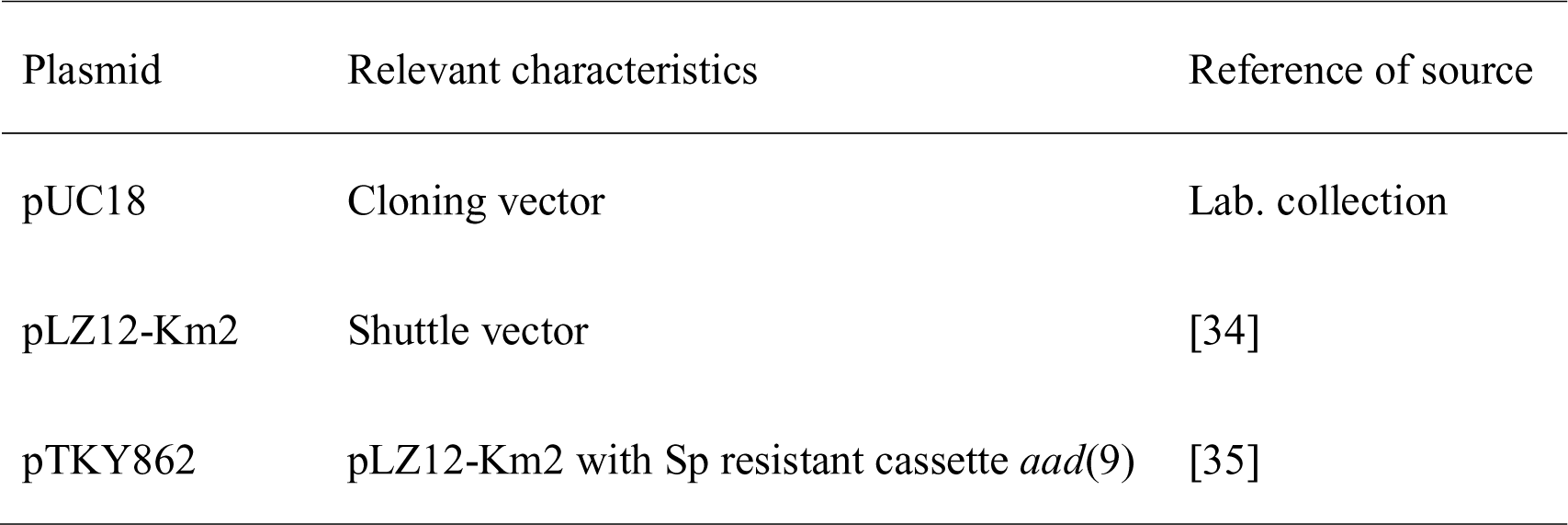
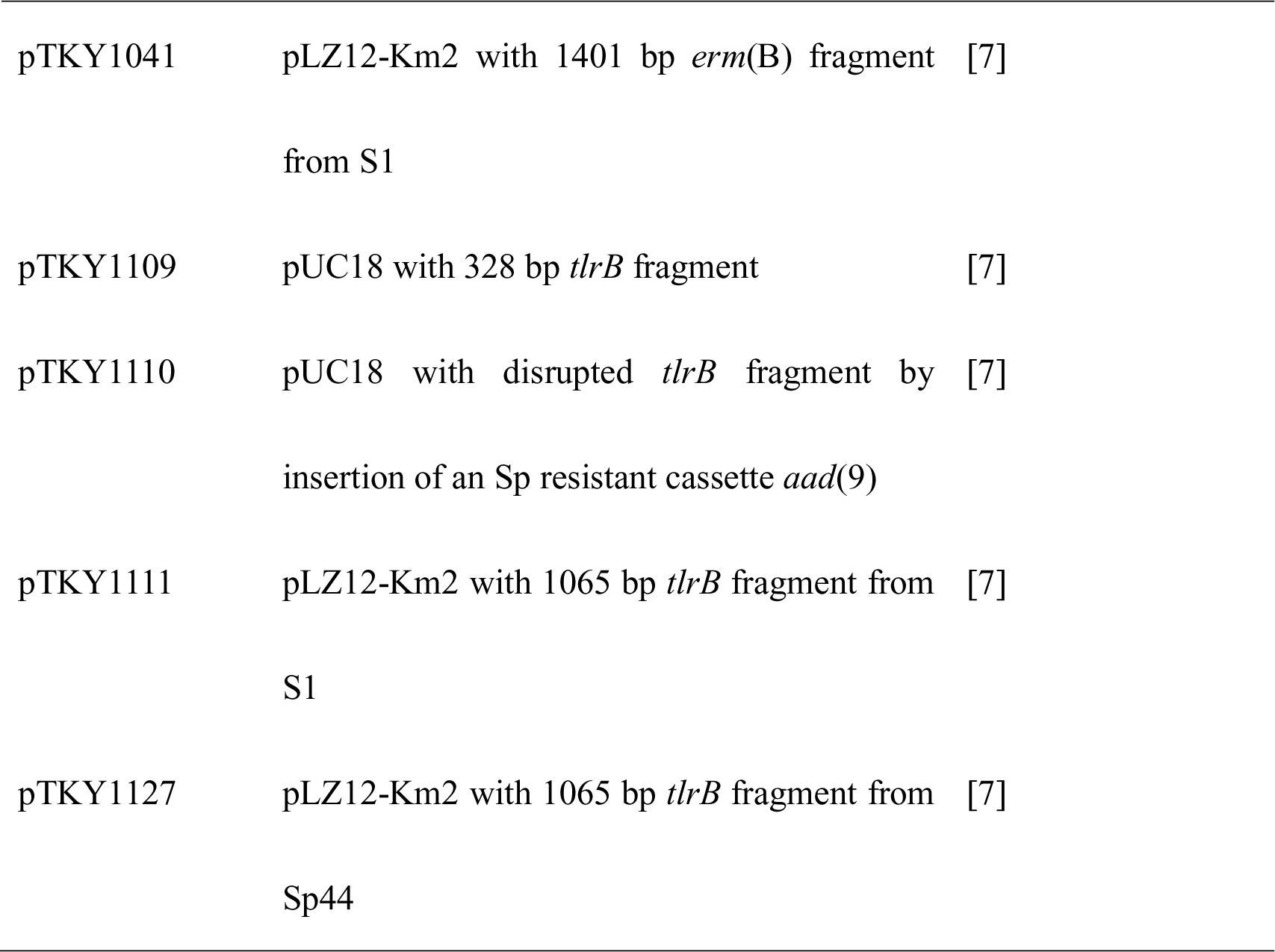
Plasmids.

Fig. 2A shows the secondary mRNA structure from the *ermBL* region that was previously demonstrated [9]. Consistent with this report, nucleotides in the stems were protected from chemical modification when probing after extracting total RNA in both Sp380 and Sp382 strains. Conversely, these regions were not protected when probing before extracting total RNA in Sp380 (Fig. 2B), but these regions remained unprotected in Sp382 (Fig. 2B). Stems were likely disrupted in Sp380 *in vivo* presumably due to ribosome stalling in the *ermBL* region. m^1^G748 affects such stalling.

**Fig 2.**
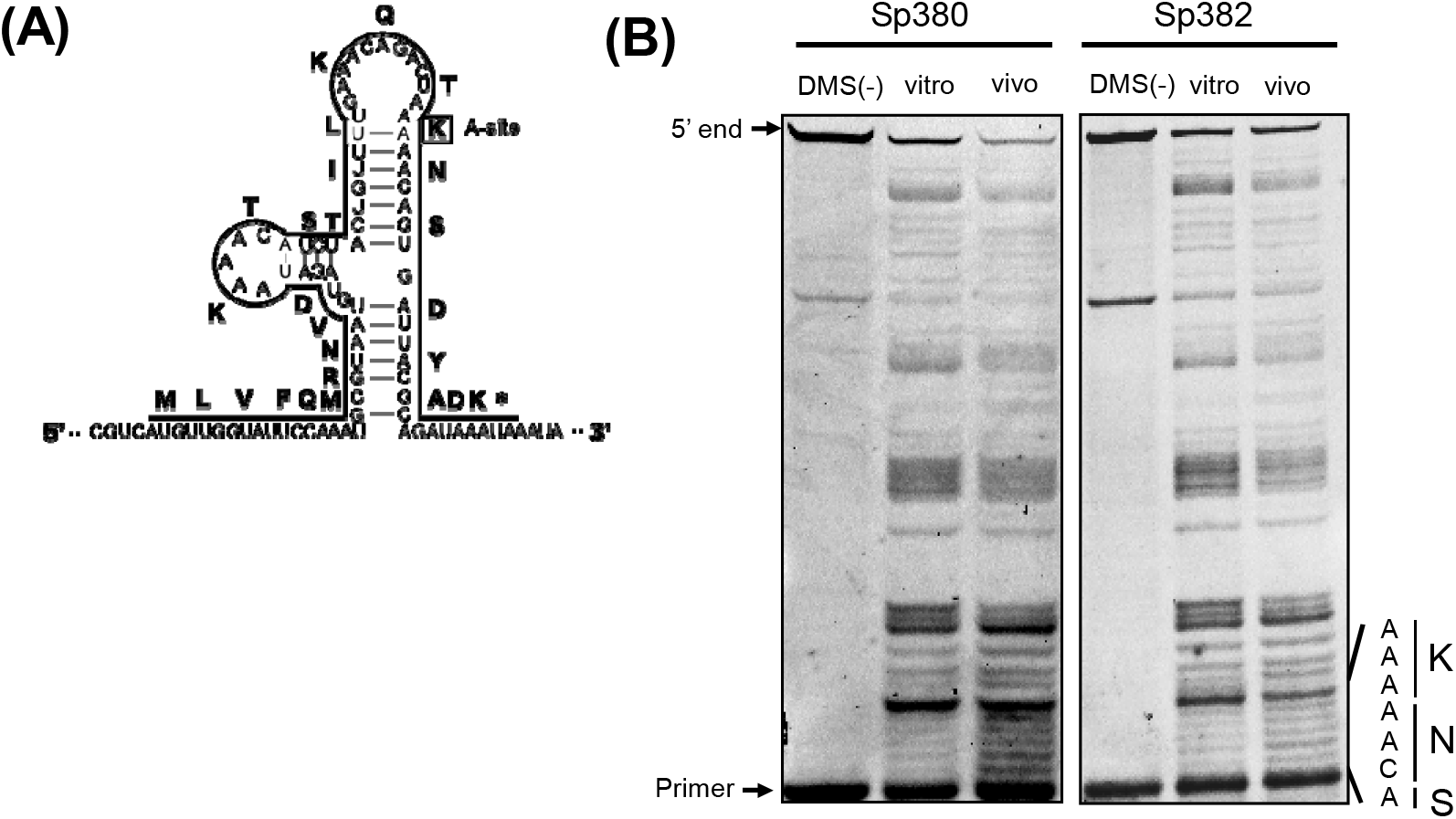
RNA footprinting assay for the *ermBL* region. (A) The secondary structure model of *ermBL* region in *erm*(B) mRNA suggested in [9]. Amino acids in at the A-site of stalled ribosomes in the *ermBL* region are boxed. (B) RNA footprinting of the *ermBL* region indicates a difference in ribosome stalling between Sp380 and Sp382. DMS was added to bacterial cultures (*in vivo*) or the total RNAs (*in vitro*). Reactions were quenched, and total RNA was purified and used in primer extension assays to detect base modifications. Unmodified RNA was used as a control (DMS (-)). The gel shows primer extension products from bases 117 to 152. The sequence around the calculated P-site and encoded amino acids are indicated on the side of the gel.

### (3) Characterization of effects of m^1^G748 on ribosome stalling

Ribosome stalling does not always mean an end to translation. Translation resumes in some cases [11, 12]. Such cases are termed “transient stalling” [13] and may contribute to cotranslational protein and be evolutionarily preferred [14, 15]. In contrast, “strong stalling” does not appear to restart and requires rescue [16, 17]. We examined the type of ribosome stalling affected by m^1^G748 by first defining A-site peaks (see Methods for details) and counted the number of A-site peaks across all CDSs (Figs. 3A and 3B). Two populations observed in Sp284 (Fig. 3A) were consistent with a previous report [16]. However, surprisingly, almost no transient stalling was observed in Sp379 (Fig. 3B), indicating a role for m^1^G748 for keeping transient stalling. A slight decrease in the number of peaks of strong stalling was observed compared to Sp284 (Figs. 3A and 3B).

**Fig 3.**
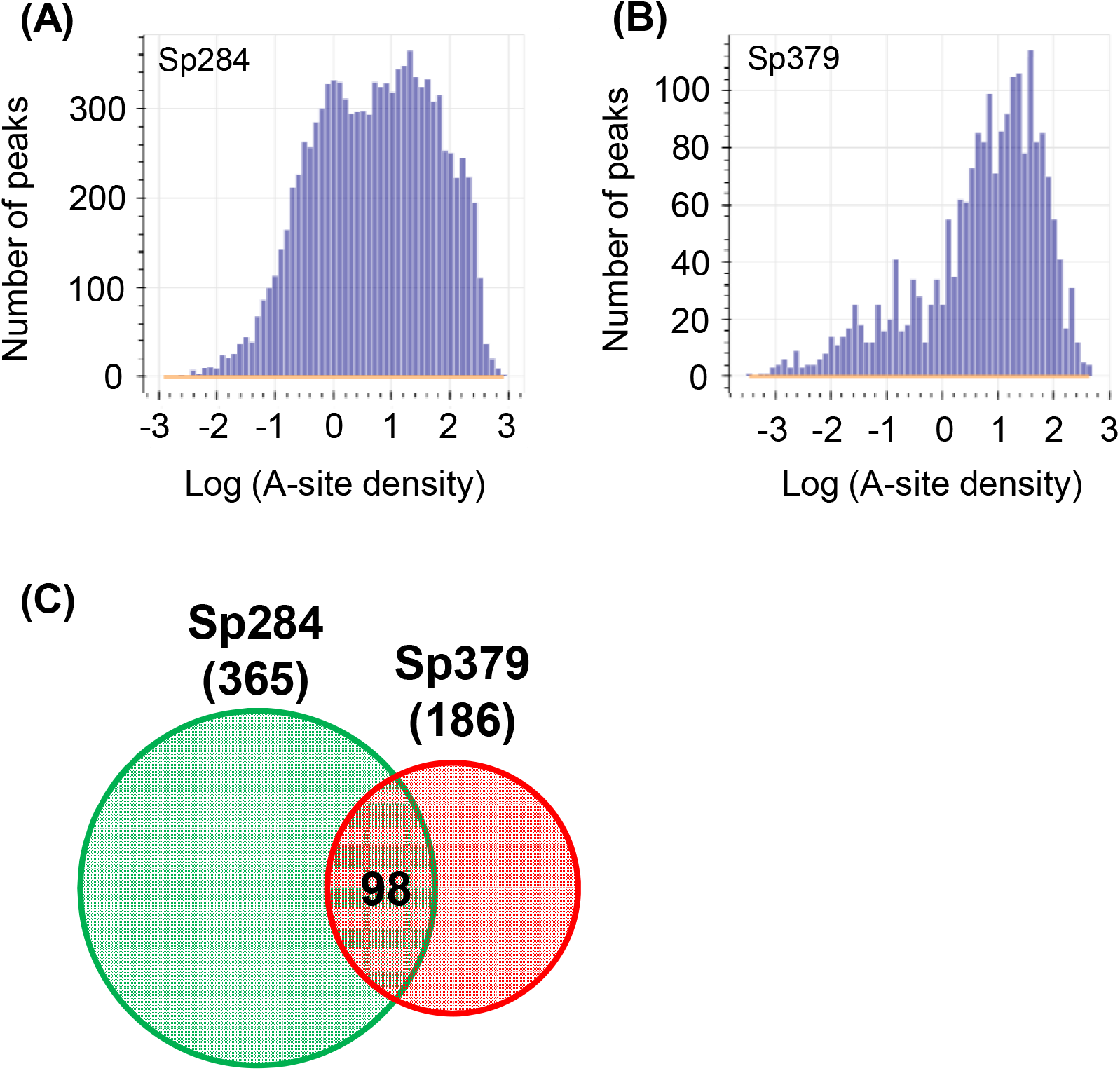
The effect of m1G748 on transient stalling and strong stalling. (A, B) Histograms showing the distribution of A-site density in Sp284 and Sp379, respectively. (C) Venn diagram for stalling peptide sequences in Sp284 and Sp379. The number of stalling peptide sequences is shown in parentheses. The number in the intersection region reflects common stalling peptides.

Stalling peptides were identified as previously described [13] to further examine the role of m^1^G748 on strong stalling. Briefly, we defined strong stalling as described in Methods and collected nascent peptide sequences in the exit tunnel for strong stalling events. We then calculated the probability of occurrence for 8,000 tripeptides and defined stalling peptides as tripeptides with a probability higher than 0.9999. Surprisingly, few stalling peptides were common between Sp284 and Sp379 (Fig. 3C), suggesting that the position of “strong stalling” is different between the two strains.

Several known stalling peptides, such as PPP, KKK and KKR, are common among some organisms, including *Saccharomyces cerevisiae* and *Escherichia coli* [18, 19]. However, stalling peptides in Sp284 and Sp379 did not include these previously reported molecules [see Additional file 1], indicating that ribosome stalling is species-specific.

### (4) m^1^G748 is required to realize evolutionarily conserved ribosome stalling

Stalling peptides in *S. pneumoniae* were different in previous reports, which led us to speculate on how they are distributed in the proteome. We examined relationships between stalling peptides and proteome in *S. pneumoniae*, by initially identifying 360 of over-represented and 382 under-represented examples for three amino acids in the *S. pneumoniae* proteome as previously [13]. Enrichment of over- and under-represented tripeptides in stalling peptides were investigated using Fisher’s exact test (Fig. 4). Over-represented peptides were significantly enriched in the set from Sp379, with under-represented peptides being significantly enriched in the stalling peptide set of Sp284, indicating that the strong stalling is not evolutionarily favored [see Additional file 2]. m1G748 is likely required to realize evolutionarily conserved ribosome stalling.

**Fig 4.**
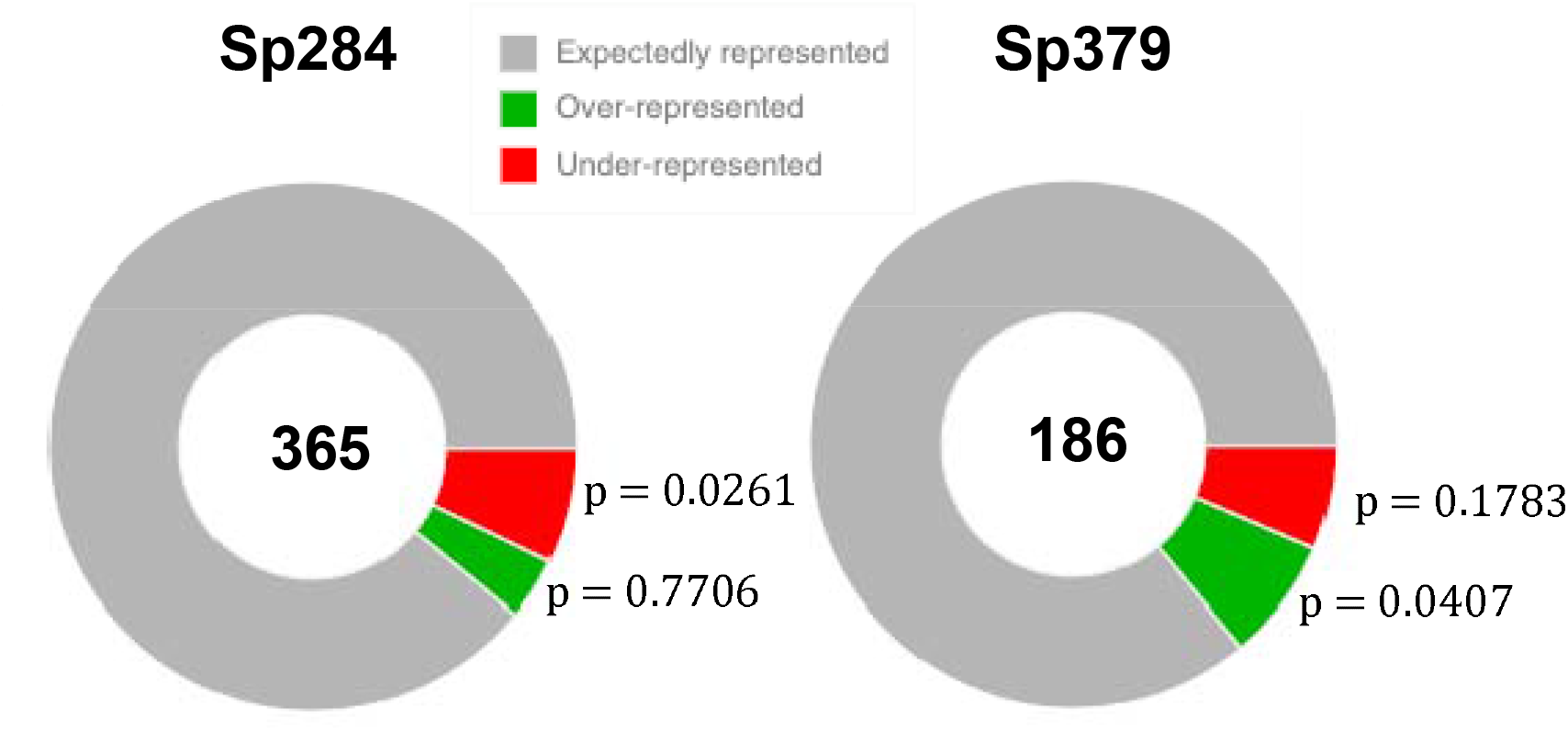
Enrichment of over- and under-represented stalling peptides. The number of the stalling peptides are represented at the center of the donut. The *p*-value represents statistical significance of the over- and under-represented fractions.

## 6. Discussion

We investigated the role of NPET in translation using a ribosome profiling assay in *S. pneumoniae*. The finding that the loss of the methyl group at m^1^G748 had a notable impact on the distribution of the ribosomes and ribosome stalling (Figs. 1 and 2) highlights the importance of the NPET structure and explains why rRNA modifications are clustered near the PTC and NPET. The role of NPET using other NPET mutants will be investigated in the future.

Alteration of NPET results in changes to the stalling peptide set (Fig. 3C). Thus, translation depends not only on mRNA sequence but also on the structure of NPET. Prediction of ribosome stalling based only on mRNA sequence would be difficult. Using NPET information is crucial because the structure of the NPET is not necessarily unique in a cell [21], with some studies having tried to predict ribosome occupancy or stalling [19,20], and performance of developed software could be improved with additional parameters for NPET structure.

A global change in stalling position may result in a global effect on cell proteome. This effect might explain why RlmA^II^ mutants of *S. pnuemoniae* are rarely clinically isolated [22]. The loss of the m^1^G748 methyl group inhibits binding of telithromycin to the exit tunnel [7], and may also cause a decrease in fitness of *S. pneumoniae* in a clinical setting. This concept might be useful since antibiotics that compromise targets that maintain healthy ribosomes would not cause resistant bacteria. TEL-resistant *S. pneumoniae* ribosomes are unhealthy.

The ribosomopathy encompasses diseases caused by abnormalities in the structure or function of ribosomal proteins or rRNA genes or other genes whose products are involved in ribosome biogenesis [23-25]. Skeletal muscle atrophy [26], Diamond–Blackfan anemia [24] and Treacher Collins syndrome [24] are examples of ribosomopathy. However, no reports regarding exit tunnel-induced ribosomopathy exist. The present study examined *S. pneumoniae*; however, exit tunnel-derived ribosomopathy might be common among organisms, including humans. For example, ribosomal protein L17 (RPL17) is upregulated in parallel with stress vulnerability [27]. RPL17 is located near the end of the exit tunnel [21]. Therefore, RPL17 could be responsible for ribosome stalling. This concept, exit-tunnel-induced ribosomopathy, might explain the mechanism of the disorder in the future.

## 7. Conclusions

We demonstrate the role of m^1^G748 on ribosome stalling in *S. pneumoniae*. m^1^G748 is required to realize species-specific and evolutionarily conserved stalling. The loss of the methyl group at m^1^G748 has a great impact 199 on the distribution of the ribosomes and ribosome stalling, and results in exit-tunnel-induced ribosomopathy. This might be the reason why RlmA^II^ mutants of *S. pnuemoniae* are rarely clinically isolated. This study is the first to show the role of NPET using an NPET mutant. These findings clarify the role of NPET structure on the translation process.

## 8. Methods

### (1) Bacterial strains, plasmids, and media

Bacterial strains and plasmids are shown in Tables 1 and 2, respectively. *S. pneumoniae* strain S1 with reduced TEL susceptibility (MIC, 2μg/ml) was clinically isolated in Japan [7]. Pneumococci were routinely cultured at 37°C and 5% CO_2_ in air in a brain-heart infusion with 0.5% yeast extract (BHI-Y) broth and BHI-Y agar, supplemented with 5% horse blood. *E. coli* was grown in L broth (1% bact-tryptone, 0.5% bact yeast extract, 0.5% sodium chloride, pH 7.4) and L agar. When necessary, medium was supplemented with kanamycin (25–500 μg/mL), spectinomycin (100 μg/mL) and ampicillin (25 μg/mL).

### (2) Transformation

Synthetic competence-stimulating peptide (CSP) 1 and the method of Iannelli and Pozzi [28] were used to transform *S. pneumoniae* S1 into a transformation-competent state.

### (3) RNA-Seq

*S. pneumoniae* cultures were grown to log-phase; 2.8 mL of cultures were added to 2.8 mL of 100°C preheated RNA lysis buffer (1% SDS, 0.1 M NaCl and 8 mM EDTA) and vortexed for 2 min. The resulting lysates were added to 5.6 mL of 100°C preheated acid phenol (Sigma-Aldrich) and vortexed for 5 min. After centrifuging, RNA was extracted from the aqueous phase using DirectZol (Zymo Research). rRNA was removed from total RNA using MICROBExpress (Ambion). Resulting total mRNA (400 ng as an input) was used for constructing the DNA library, using KAPA Stranded RNA-Seq Library Preparation Kit Illumina platforms (KK8400). DNA libraries were sequenced using the Illumina HiSeq 1500 system and single-end reads. Illumina libraries were preprocessed by clipping the Illumina adapter sequence using Trimmomatic v.0.39 [29], then aligned to the S1 genome sequence [7] using HISAT2 v.2.2.1 [30].

### (4) Ribo-Seq

Libraries were prepared as previously described with some modifications (see below) [10].

### Cell growth and harvest

*S. pneumoniae* cultures (2.4 L) were grown to log-phase. Cells were pretreated for 2 min with ∼ 100 μg/mL chloramphenicol. Immediately after chloramphenicol pretreatment, cultures were placed on ice. Cells were pelleted by centrifugation at 8,000 × *g* for 15 min at 4°C. After decanting the supernatant, cell pellets were resuspended in 2.5 mL of prechilled resuspension buffer [10 mM MgCl_2_, 100 mM NH_4_Cl, 20 mM Tris (pH 8.0), and 1 mM chloramphenicol].

### Lysate preparation and Nuclease digestion

Cells were sonicated on ice and centrifuged at 12,000 × *g* for 10 min at 4°C. Aliquots of lysate, containing 25 Abs_260_ ribosome units (1 A_260_ = 12 μg/μL) [5] were digested with 60 U of MNase (Roche) and 60 U of SUPERase·In (Ambion) with the addition of chloramphenicol to a final concentration of 1 mM to remove unprotected mRNA and generate footprint fragments. Digestion reactions were incubated for 1 h at 25°C and quenched with the addition of EGTA to a final concentration of 6 mM.

### Sucrose fractionation

Linear sucrose gradients [5–40% (wt/vol)] were prepared with 7.6 mL of buffer A and buffer B [10 mM MgCl_2_, 100 mM NH_4_Cl, 2 mM DTT, 20 mM, 0.2 mM chloramphenicol, Tris pH 7.8 and 5% or 40% Sucrose, respectively], by loading Buffer B on Buffer A in 16PA tube and placed vertically for 12 hr at 4°C the placed horizontally for 2 hr at 4°C.

Digested samples were carefully loaded onto prepared gradients and centrifuged at 124,700 × *g* for 8 hr at 4°C in a P28S2 rotor. Sucrose gradients were fractionated manually into 200 μL portions and A_260_ was recorded for each fraction. Absorbance values were graphed in Microsoft Excel to determine ribosome footprint associated fractions (RPF fractions). RPF were consequently pooled.

### Ribosome footprint preparation

RNA was purified from RPF using the SDS/hot acid phenol. Three mL of samples were first denatured with SDS to a final concentration of 1% (wt/vol) and 2.7 mL of preheated acid phenol (65°C) (Sigma-Aldrich). Mixtures were vortexed for 5 min at 65°C. After centrifuging, aqueous phases were mixed with one volume of acid phenol, and 0.9 volumes of chloroform/isoamyl alcohol (24:1). RNA was precipitated with ethanol and resuspended in 10 μL of 10 mM Tris (pH 8.0).

Ribosome footprint samples were resolved on denaturing polyacrylamide gels for size selection of footprint fragments. RNA samples were prepared for electrophoresis by adding 2x Novex TBE-Urea Sample Buffer (Invitrogen). Ladder standards used a 0.05 μg/μL 10-bp DNA Ladder (Invitrogen), prepared in 2x Novex TBE-Urea Sample Buffer and 10 mM Tris (pH 8.0). Samples were resolved on a 15% TBE-Urea gel in 1 × TBE buffer for 65 min at 200V. Gels were stained for 3 min with SYBR Gold Nucleic Acid Gel Stain (diluted from 10,000x in 1x TE; Invitrogen) and visualized by UV transillumination. A band between 20 and 45 was excised using the 10-bp DNA ladder to identify footprint fragments. RNA was recovered using the ZR small-RNA PAGE Recovery Kit (Zymo Research) following the manufacturer’s protocol, except that RNA was eluted from the final spin column with 15 μL of 10 mM Tris (pH 8.0). Collected RNA was quantified and characterized by using a small-RNA chip on the Agilent BioAnalyzer (Agilent Technologies).

### Dephosphorylation

Three hundred pmol of footprint were denatured for 2 min at 80°C and placed on ice. The 3’ ends were dephosphorylated with T4 polynucleotide kinase (T4 PNK; NEB) in the following reaction mix: 1 × T4 PNK reaction buffer (without ATP), 20 U SUPERase·In, and 10 U T4 PNK. Reactions were incubated at 37°C for 1 hr. The enzyme was then heat-inactivated for 10 min at 75°C. RNA was precipitated with isopropanol and resuspended in 10 μL of 10 mM Tris (pH 8.0).

### Linker-1 Ligation

Twenty pmol of dephosphorylated RNA was prepared by diluting with 10 mM Tris (pH 8.0). One μg of Linker-1 (5’-App CTGTAGGCACCATCAAT ddC-d’) was added to RNA samples. Mixtures were denatured for 90 s at 80°C, then cooled to room temperature for 15 min. Ligation of RNA to Linker-1 used the following reaction components: 20% (wt/col) PEG, 10% DMSO, 1 × T4 Ligase reaction buffer, 20 U SUPERase · In, and 10 U T4 Ligase 2, truncated (NEB). Reaction mixtures were incubated at 37°C for 1 h. 2 × TBE-Urea sample buffer was added to reaction mixtures. Samples were resolved on a 10% TBE-Urea gel in 1 × TBE buffer at 200V. Gels were stained for 3 min in SYBR Gold Nucleic Acid Gel Stain and visualized by UV transillumination. A band between 30 and 70 nt was excised using the 10-bp DNA ladder to mark ligated product size. Ligated RNA was recovered using the ZR small-RNA PAGE Recovery kit. Ligated products were eluted from the final spin column with 6 μL of 10 mM Tris (pH 8.0).

### Phosphorylation

The collected 3’ ligated samples were incubated for 2 min at 80°C and placed on ice. The 5’ ends were phosphorylated by T4 PNK in the following reaction mix: 1 × T4 PNK Reaction Buffer, 20 U SUPERase·In, 10 U T4 PNK and 1 mM ATP. Reactions were incubated at 37°C for 1 hr. The enzyme was heat-inactivated for 10 min at 75°C. RNA was precipitated with isopropanol and resuspended in 6 μL of 10 mM Tris (pH 8.0).

### Linker-2 ligation

1 μL of 100 μM Linker-2 (5’-GAGTCTGCGTGTGATTCGGGTTAGGTGTTGGGTTGGGCCA-3’) was added to phosphorylated RNA samples. Mixtures were denatured for 15 min at 65°C and placed on ice. Ligation of RNA to Linker-2 used the following reaction mix: 17.5% (wt/vol) PEG, 1 × T4 ligase reaction buffer, 20 U SUPERase·In, and 10 U T4 RNA Ligase1 (NEB). Reaction mixtures were incubated at 37°C for 2.5 h. Two × TBE-Urea sample buffer was added and ligated RNA resolved on 10% TBE-Urea gels in 1 × TBE buffer at 200V. Gels were stained for 3 min in SYBR Gold Nucleic Acid Gel Stain and visualized by UV transillumination. A band between 90 and 120 nt was excised using the 10-bp DNA ladder to identify ligated products. Ligated RNA was recovered using the ZR small-RNA PAGE Recovery kit. Products were eluted from the final spin column with 6 μL of 10 mM Tris (pH 8.0).

### Reverse transcription

Four point five μL of ligated samples were mixed with 1 μL of 0.1 μM Linker-1-RT (5’-ATTGATGGTGCCTACAG-3’) and 1 μL of 0.5 mM dNTP. The resulting mixtures were denatured for 2 min at 80°C then quickly cooled on ice. Samples were incubated at room temperature for 10 min reverse transcription using SuperScript III Reverse Transcriptase (Invitrogen) with the following reaction mix: 1 × first strand buffer, 5 mM DTT, 20 U SUPERase·In and 200 U SuperScript III Reverse Transcriptase. Reaction mixtures (10 μL) were incubated for 1 hr at 47°C. RNA products were hydrolyzed by adding 1 mM NaOH to a final concentration of 0.1 mM and incubated for 15 min at 95°C. cDNA products were resolved from the unextended primer on a 10% TBE-Urea gel in 1 × TBE buffer at 200V. Samples were prepared for electrophoresis by adding 2 × TBE-Urea sample buffer and denaturing for 5 min at 95°C. Gels were stained for 3 min in SYBR Gold Nucleic Acid Gel Stain and visualized by UV transillumination. A band between 90 and 120 nt was excised using the 10-bp DNA ladder to identify reverse transcription products. DNA was recovered using the ZR small-RNA PAGE Recovery Kit. cDNA products were eluted from the final spin column with 6 μL of 10 mM Tris (pH 8.0).

### 2nd strand DNA synthesis

cDNA (100 pg) was amplified with Q5 High Fidelity Polymerase (NEB) using LInker-2-partial (5’-TTAGGTGTTGGGTTGGGCCA-3’) and Linker-1 as primers. Amplified PCR products were purified using AMPure Bead (Beackman Coulter). Double-stranded DNA was eluted from beads with 10 μL of 10 mM Tris (pH 8.0).

### Library preparation and sequencing

KAPA Hyper Prep Kits from Illumina (KK8500) were used to construct the library with a slight modification to the manufacturer’s protocol. The manufacturer’s protocol followed the EndRepair and A-tailing steps, where, briefly, 5 μL of KAPA frag buffer was added to 45 μL of purified double-stranded DNA. The resulting DNA library was quantified and characterized using the high-sensitivity DNA chip on an Agilent BioAnalyzer (Agilent Technologies). Libraries were sequenced using an Illumina HiSeq 1500 system and single-end reads after adding PhiX (Illumina) to a final concentration of 30% (vol/vol) to improve sequencing quality.

### Sequence analysis

Illumina libraries were preprocessed by clipping adapter sequence (Linker-1 and Linker-2) using TrimmomaticPE v.0.39 [29] for the Illumina adapters and CutAdapt v.2.10 [31] for linker sequences. Sequencing reads were aligned to the S1 genome sequence [7] using HISAT2 v.2.2.1 [30]. The S1 gene feature file [7] was used to identify the CDS region. Sequencing data were deposited in the DDBJ database with accession number DRA011224.

### (5) Definition of the ribosome density

The ribosome density (RD) was calculated as previously described [32], except that ribosome footprints between 24 and 30 nt were selected for calculations. This range was chosen to assess as many footprints as possible to improve statistical power and to exclude fragments suspected not to represent footprints (see Additional file 3 for the detail).

### (6) Metagene analysis

Each normalized RD profile was aligned by its start codon and averaged across each position, with RD profiles first being scaled by their own mean density to obtain metagene profiles (Fig. 1B).

### (7) DMS modification of ribosomes

Dimethyl sulfate (DMS) modification was based on a previously published protocol [7]. *S. pneumoniae* cultures (2.4 L) were grown to log-phase. Cells were pelleted by centrifugation at 8,000 × *g* for 15 min at 4°C and washed and resuspended in 250 μL of DMS buffer [10 mM MgCl_2_, 50 mM sodium cacodylate pH 7.0]. For the modification *in vivo*, DMS solution was added to a final concentration of 50 mM and incubated for 1 hr at 20°C. The reaction was quenched by the addition of 62.5 μL ice-cold stop buffer [1 M β-mercaptoethanol and 1.5 M sodium acetate, pH 7.0]. RNA was extracted using SDS/hot acid phenol method. Preheated (100°C) acid phenol (0.9 volumes) was added to the quenched mixture and vortexed for 5 min. After centrifugation, RNA was purified from the aqueous phase using DirectZol (Zymo Research). For the modification *in vitro*, RNA was first extracted from the cell suspension using the same method as for *in vivo*. After purifying the total RNA, DMS solution was added to a final concentration of 50 mM in DMS buffer and incubated for 1 hr at 20°C then quenched. RNA was precipitated with ethanol and resuspended in 20 μL of 10 mM Tris (pH 8.0).

### (8) Primer extension

The degree of methylation of each RNA was assayed by primer extension initially described by Morgan et al. [33]. Briefly, 210 μg of DMS-probed total RNA, 400 Nm concentration of an oligonucleotide primer containing 5’ - linked fluorescein isothiocyanate (FITC) (5’-[FITC]GAATTAATCTAACGTATTTATTTATCTGCGTAATCA -3’), and 500 μM dNTP were mixed, heated to 95°C for 1 min, and cooled to allow annealing at 53°C. A mixture containing SuperScript III Reverse Transcriptase was added, and the reaction was continued at 53°C for an additional 20 min. Reactions were terminated by the addition of 2 × TBE-Urea sample buffer after incubation at 70°C for 15 min. Extension products were resolved on a 12.5% TBE-urea gel and visualized using a TyphoonFLA9000 photoimager (GE Healthcare).

### (9) Definition of A-site peaks

We first determined the A-site corresponding to each read by an offset of [(15/27)×(L)] from the 5’ end of the read, where L is the length of each read. Then we used normalized A-site count (ribo/mRNA) as A-site peaks.

### (10) Definition of strong stalling

Strong stalling is defined as A-site peaks with a height of higher than eight because the number of A-site peaks in the *ermBL* region was around eight. This region is where ribosome stalling was confirmed by RNA footprinting. The results, however, remain robust also for other cutoff values of A-site density.

### (11) Enrichment of over-, under-represented peptide sequence

Over- and under-represented peptide sequences were identified as previously described [13], except that *p*-value cutoff is 0.0001 for under-represented peptides and 0.9999 for over-represented peptides.

## 10. Supplementary Information

Additional file 1: Supplementary Tables S1

Additional file 2: Supplementary Tables S2

Additional file 3: Supplementary Fig. S1

## 11. Declarations

### (1) Ethics approval and consent to participate

Not applicable.

### (2) Consent for publication

Not applicable.

### (3) Availability of data and materials

All data generated in this study have been deposited to the DDBJ depository (DRA011224).

### (6) Author contributions

ST contributed to the conception and overall design of the work, all of experiments, bioinformatics analysis and interpretation of Ribo-Seq data and drafting of the manuscript. TA contributed to the design of Ribo-Seq experiments. KY contributed to Ribo-Seq experiments. TH contributed to bioinformatics analysis, interpretation of Ribo-Seq data, drafting of the manuscript and substantial revision of the manuscript. YT contributed to the drafting of the manuscript. KH contributed to the conception and overall design of the work, interpretation of data, and drafting and revision the manuscript. The authors read and approved the final manuscript.

## Acknowledgments

We thank Tomoko Yamamoto for discussions and comments to the manuscript.

## 14. Supplementary Figure Legends

**Fig. S1 Histograms showing the distribution of length of RFPs in Sp284 and Sp379, respectively**.

## 15. Supplementary Table Legends

**Table S1 The list of stalling peptides**.

The first and the second columns list stalling peptides in Sp284 and Sp379 respectively. The third column shows the list of stalling peptides that are common between strains.

**Table S2 The list of over- and under-represented peptides**.

The first and the second columns list over-represented peptides and under-represented peptides in *S. pneumoniae* proteome composition.

