## Supplementary figures and images for "Ribosome profiling in *Streptococcus pneumoniae* reveals the role of methylation of 23S rRNA nucleotide G748 on ribosome stalling"

### Additional file 3

#### Slide 1
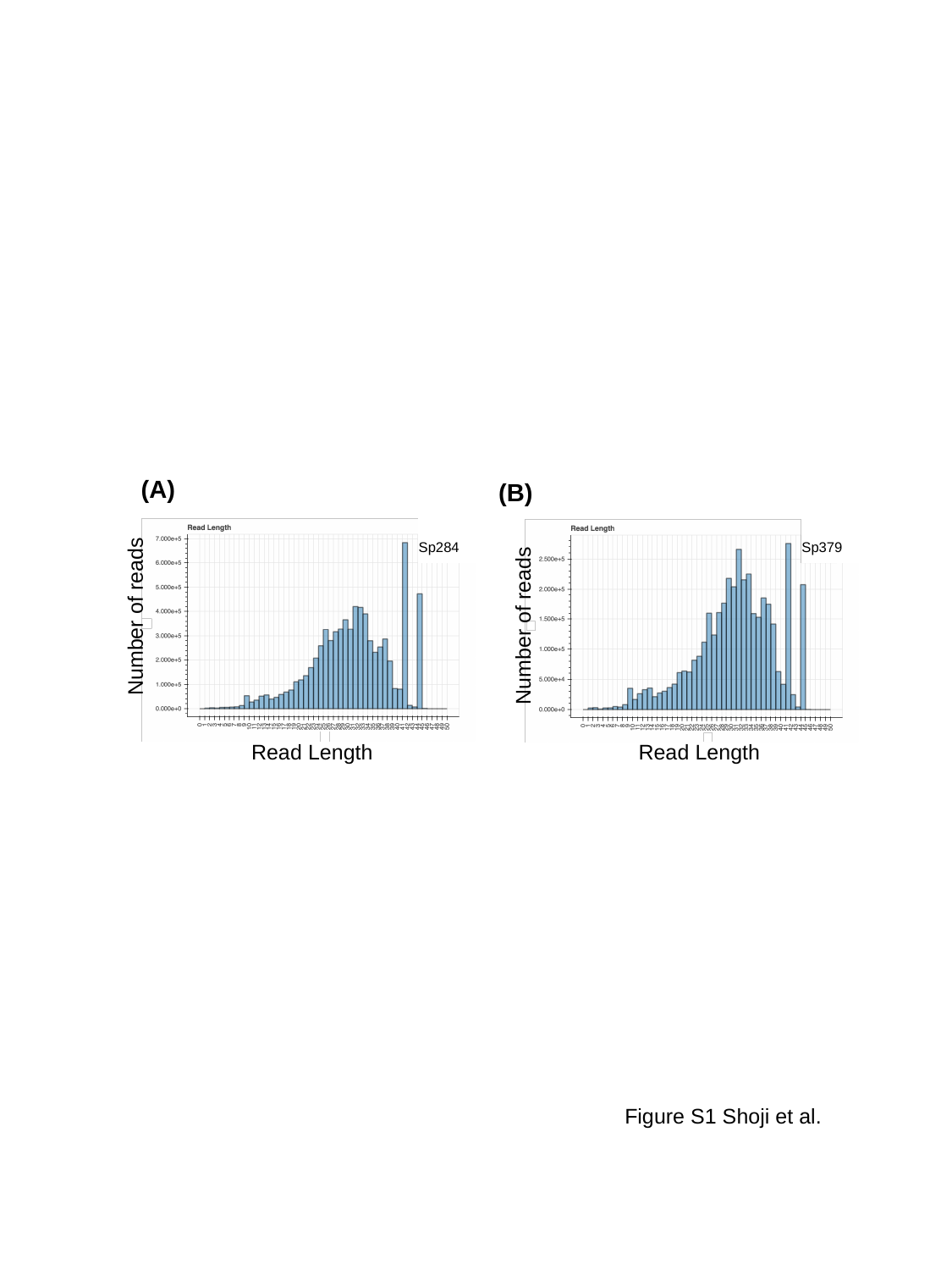

(A)
(B)
Sp284
Number of reads
Read Length
Sp379
Number of reads
Read Length
Figure S1 Shoji et al.
